# An anti-CRISPR protein induces strong non-specific DNA binding activity in a CRISPR-Cas complex

**DOI:** 10.1101/2020.05.28.119941

**Authors:** Wang-Ting Lu, Chantel N. Trost, Hanna Müller-Esparza, Lennart Randau, Alan R. Davidson

**Affiliations:** Department of Biochemistry, University of Toronto, Toronto, Canada; Department of Computer Science and Engineering, National Institute of Technology, Hamirpur, Himachal Pradesh, India; Faculty of Biology, University of Marburg, Germany; Loewe Center for Synthetic Microbiology (SYNMIKRO), Germany

**Keywords:** CRISPR-Cas, anti-CRISPR, non-specific DNA binding, phage

## Abstract

Phages and other mobile genetic elements express anti-CRISPR proteins (Acrs) to protect their genomes from destruction by CRISPR-Cas systems. Acrs usually block the ability of CRISPR-Cas systems to bind or cleave their nucleic acid substrates. Here, we investigate an unusual Acr, AcrIF9, that induces a gain-of-function to a type I-F CRISPR-Cas (Csy) complex, causing it to bind strongly to DNA that lacks both a PAM sequence and sequence complementarity. We show that specific and non-specific dsDNA compete for the same site on the Csy:AcrIF9 complex with rapid exchange, but specific ssDNA appears to still bind through complemetarity to the CRISPR RNA. We also demonstrate that induction of non-specific DNA-binding is a conserved property of diverse AcrIF9 homologues, implying that this activity contributes the biologically relevant function of this Acr family. AcrIF9 provides another example of the surprising variety of mechanisms by which Acrs inhibit CRISPR-Cas systems.

## INTRODUCTION

<u>C</u>lustered <u>R</u>egularly <u>I</u>nterspaced <u>S</u>hort <u>P</u>alindromic <u>R</u>epeats (CRISPR) and CRISPR-associated (Cas) proteins together represent an adaptive mechanism employed by many species of bacteria and archaea to destroy potentially harmful mobile genetic elements (MGEs), such as phages (Bondy-Denomy & Davidson, 2014; Makarova et al., 2020; Makarova, Wolf, & Koonin, 2018). CRISPR arrays are comprised of repeated DNA sequences interspersed with non-repeating DNA spacer sequences. Spacer DNA is often identical to MGE sequences. These arrays are transcribed and processed into CRISPR (cr) RNAs, comprised of one repeat and one spacer, that form complexes with Cas proteins. Using the spacer sequence as a guide, CRISPR-Cas complexes specifically bind DNA or RNA and mediate subsequent nucleolytic destruction of the targeted nucleic acid. CRISPR-Cas systems provide adaptive immunity against foreign DNA as segments of newly encountered MGEs can be incorporated into CRISPR arrays in the form of new spacers, providing defence against subsequent encounters with the same MGE.

CRISPR-Cas systems are tremendously diverse with 33 distinct subtypes distributed among 6 different types (Makarova et al., 2020). Here, we focus on the type I-F system, which is found widely in *Proteobacteria.* The type I-F CRISPR-Cas complex, which is known as the Csy complex, comprises a 60 nucleotide (nt) crRNA with one molecule of Cas6f bound to the 3’-hairpin formed by the repeat sequence followed by 6 molecules of Cas7f bound to the 32 nt spacer (van Duijn et al., 2012; Wiedenheft et al., 2011). A complex of Cas5f and Cas8f are bound to another portion of the repeat sequence known as the handle, which lies at the 5’-end of the crRNA (Chowdhury et al., 2017; Guo et al., 2017; Rollins et al., 2019). Binding of dsDNA targets is initiated by Cas8f recognition of the protospacer adjacent motif (PAM) and subsequent separation of the DNA strands. Known as R-loop formation, this strand separation allows for hydrogen-bonding between the spacer region of the crRNA and the target strand of the DNA. Large conformation changes in the Csy complex occurring upon target DNA-binding lead to exposure of a Cas8f domain that recruits Cas3, the helicase-nuclease that mediates processive degradation of the targeted DNA (Rollins et al., 2019).

Our group discovered the first phage-encoded inhibitors of a CRISPR-Cas system, describing five families of anti-CRISPR proteins (Acrs) that blocked the activity of the type I-F system of *Pseudomonas aeruginosa (Pae)* (Bondy-Denomy, Pawluk, Maxwell, & Davidson, 2013). Since then, more than 60 families of Acrs have been described that act against many different types of CRISPR-Cas systems (Bondy-Denomy et al., 2018). Mechanistic and structural studies on Acrs have provided new insights into how CRISPR-Cas systems function and have illustrated the many fascinating ways by which small proteins can inhibit large protein-RNA complexes (Athukoralage et al., 2020; Bondy-Denomy et al., 2015; Davidson et al., 2020; Dong et al., 2019; Guo et al., 2017; Harrington et al., 2017; Wang et al., 2016). In addition, a growing number of biotechnological applications for Acrs are being developed (Marino, Pinilla-Redondo, Csorgo, & Bondy-Denomy, 2020; Nakamura et al., 2019; F. Zhang, Song, & Tian, 2019).

Despite numerous studies on Acr mechanisms, detailed biochemical and structural investigations have been carried out on only three Acrs specific to the I-F system (Bondy-Denomy et al., 2015; Chowdhury et al., 2017; Guo et al., 2017; Wang et al., 2016). Since these characterized Acrs each function through distinct mechanisms, we reasoned that the study of additional I-F Acrs would reveal new means of inhibition. To this end, we chose to investigate the AcrIF9 family, which is one of the largest and most diverse families of I-F Acrs (Pawluk et al., 2016). In a recent structural study on the Csy complex bound to AcrIF9 (Csy:F9), we and our collaborators found that this complex binds to dsDNA in a non-specific manner, requiring neither a PAM nor complementarity to the crRNA (Hirschi et al., 2020). Here, we have utilized a variety of biochemical approaches to characterize and understand this surprising property of the Csy:F9 complex. This work corroborates the structural studies previously performed on AcrIF9 (Hirschi et al., 2020; K. Zhang et al., 2020), and illuminates the unique features of this anti-CRISPR.

## RESULTS

### AcrIF9 binds to Cas7f at a site overlapping with AcrIFI

Genes encoding four members of the AcrIF9 family found in strains of *Vibrio parahaemolyticus* (*Vpa*), *Proteuspenneri* (*Ppe\ Aggregatibacter actinomycetemcomitans* (*Aac*), and *Xanthomonas fragariae* (*Xfr*) were synthesized and expressed in *Pae* (Fig. S1*A*). All four homologs showed robust inhibition of the *Pae* type I-F CRISPR-Cas system (Fig. S1*B*). Expression of 6xHis-tagged versions of each homolog in *E. coli* followed by Ni-NTA purification showed that the homolog from *Ppe* was most suitable for further biochemical analysis due to its displaying the highest expression level and solubility. All subsequent studies were performed with this protein except where noted.

Two different recently solved cryo-EM structures of AcrIF9-bound Csy (Csy:F9) complex revealed a stoichiometry of two AcrIF9 monomers per complex (Hirschi et al., 2020; K. Zhang et al., 2020). Remarkably, the binding sites of AcrIF9 overlap very closely with those of the previously characterized anti-CRISPR, AcrIF1 (Bondy-Denomy et al., 2015; Chowdhury et al., 2017). To confirm that AcrIF9 and AcrIF1 occupy overlapping sites on Cas7f, we conducted a competitive binding experiment. Untagged AcrIF9 was added to Csy complex containing 6xHis-tagged Cas7f that was pre-saturated with untagged AcrIF1. Purification of Acr-bound Csy complexes using Ni-NTA chromatography showed that the prior addition of AcrIF1 completely blocked the binding AcrIF9 (Fig. S2). By contrast, prior addition of AcrIF9 or AcrIF1 to the Csy complex did not impede the binding of untagged AcrIF2, which binds to Cas8f (Bondy-Denomy et al., 2015; Chowdhury et al., 2017). These results confirm that AcrIF9 and AcrIF1 bind to the same binding site on Cas7f.

### AcrIF9 abolishes the DNA binding specificity of the Csy complex

Since AcrIF9 and AcrIF1 share binding sites on the Csy complex, it was expected that they would have the same effect on the Csy complex. AcrIF1 sterically blocks the hybridization of target DNA to the crRNA, thus strongly inhibiting the binding of both dsDNA and ssDNA (Guo et al., 2017). However, previous work indicated that AcrIF9 induced the Csy complex to bind DNA non-specifically, a property not observed for AcrIF1 (Hirschi et al., 2020). To investigate this surprising aspect of AcrIF9, Electrophoretic Mobility Shift Assays (EMSAs) were used to assess the effect of AcrIF9 on the DNA-binding activity of the Csy complex. As shown previously, binding of the Csy complex to a 50 bp dsDNA fragment containing a sequence matching the crRNA spacer and an appropriate Protospacer Adjacent Motif (PAM) causes a large change in the mobility of the fragment in a polyacrylamide gel ((Bondy-Denomy et al., 2015), Fig. 1*A*). Addition of AcrIF9 to the Csy complex did not abrogate dsDNA binding, and the shifted band displayed a slower mobility and was smeared (Fig. 1 *A*). By contrast, a Csy complex bound to AcrIF1 (Csy:F1) displayed no dsDNA-binding ability, as was previously shown (Bondy-Denomy et al., 2015). When specific ssDNA was used as a binding substrate, the Csy:F9 complex bound as well as the Csy complex alone; again contrasting with the Csy:F1 complex that displayed greatly reduced ssDNA binding (Fig. 1 *A*). Thus, AcrIF9 and AcrIF1 elicit very different effects on the DNA-binding activity of the Csy complex despite binding at overlapping sites on Cas7f.

**Fig. 1.**
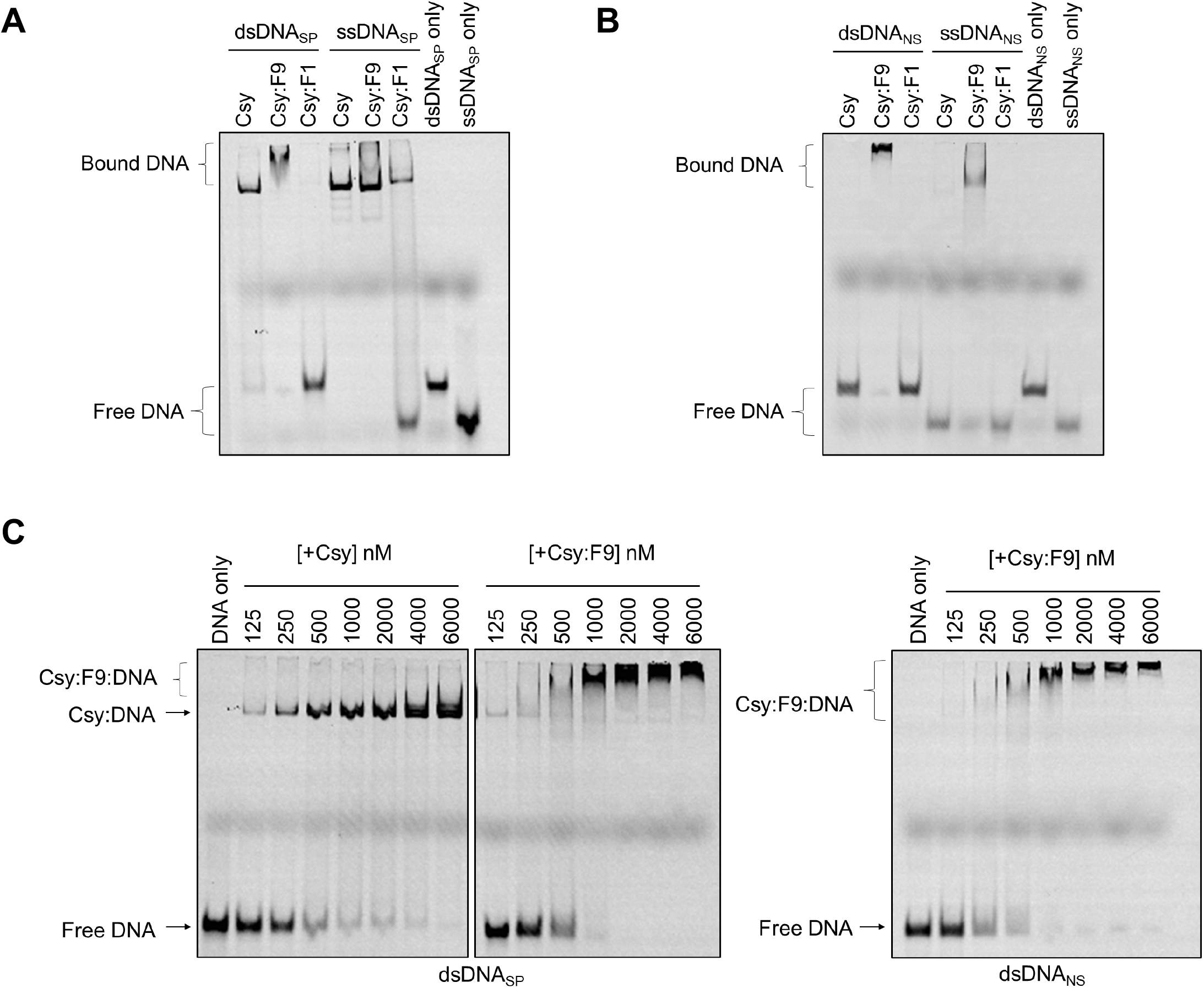
The Csy:F9 complex binds to DNA non-specifically. (*A*) The binding of Csy, Csy:F9, and Csy:F1 complexes to dsDNA_SP_ and ssDNA_SP_ was assessed on EMSA gels. (*B*) The same experiment shown in (*A*) was performed with dsDNA_NS_ and ssDNA_NS_. (*C*) Increasing concentrations of Csy or Csy:F9 complex were added to constant concentrations of dsDNA_SP_ (left two panels) or dsDNA_NS_ (right panel). These experiments were performed using 50 bp DNA fragments 5’-labeled with Cyanine 5 (Cy5).

The smearing of the dsDNA band bound to Csy:F9 suggested that this complex may not recognize a specific site on the DNA. To test this idea, we performed a similar EMSA experiment using a 50 bp non-specific dsDNA sequence (dsDNA_NS_) that possesses the same base composition as the specific target DNA, but the sequence is randomized. The PAM sequence is also absent. While the Csy complex on its own displayed no affinity for dsDNA_NS_, Csy:F9 displayed robust binding activity (Fig. 1*B*). Notably, binding of Csy:F9 to dsDNA_NS_ resulted in the a similar supershifted and smeared band as was seen in the experiments with the specific target DNA (dsDNA_SP_). Csy:F9 also bound to non-specific ssDNA (ssDNA_NS_). AcrIF9 alone showed no binding to dsDNA (Fig. S3). Titration experiments where concentrations of unbound Csy or Csy:F9 were incrementally increased indicated that the affinity of Csy:F9 for dsDNA_NS_ was similar to its affinity for dsDNA_SP_ and to the affinity of unbound Csy for dsDNA_SP_. In each of these cases, the DNA was mostly bound at a complex concentration of 1000 nM (Fig. 1*C*). It should be noted that the EMSA results involving AcrIF9 presented here differ in appearance from those previously published (Hirschi et al., 2020). These previous assays were run under different conditions. In addition, these assays used a constant concentration of both DNA and Csy complex, only increasing the concentration of AcrIF9. Here, the Csy:F9 complexed was preformed through co-expression and purified as a complex. Increasing concentrations of the preformed complex were added to the reactions.

### Multiple Csy:F9 complexes bind to non-specific DNA

To further investigate the DNA-binding properties of Csy:F9, we used a fluorescence polarization (FP) assay (Anderson, Larkin, Guja, & Schildbach, 2008). The binding of fluorescently labeled DNA to the 350 kD Csy complex markedly reduces its tumbling rate causing an increased FP signal. Thus, the binding of DNA to Csy and Csy:F9 could be quantitated by monitoring the FP signal. Consistent with the EMSA results, the binding of Csy:F9 to both dsDNA_NS_ and dsDNA_SP_ was readily detected and occurred at Csy complex concentrations within the same range as required for binding of dsDNA_SP_ to the Csy complex on its own (Fig. 2*A*). Dissociation constants *(Kd* values) calculated from these data showed the Csy complex binding dsDNA_SP_ with a *K*_d_ of 17 ± 6 nM while Csy:F9 bound dsDNA_SP_ and dsDNA_NS_ with apparent *K*_d_ values of 96 ± 33 nM and 73 ± 23 nM, respectively. These values were calculated assuming formation of a 1:1 Csy:DNA complex, which is likely not the case for Csy:F9 as is discussed below; thus, we use the term “apparent *K*_d_” and quote these values only to provide an estimate of the binding strength. Notably, the FP signal measured for the Csy:F9 complex binding to dsDNA_NS_ at saturation was nearly double that seen when dsDNA_SP_ was bound by Csy alone, suggesting that more than one molecule of Csy:F9 may be binding to each molecule of dsDNA_NS_. Supporting this idea, the Csy:F9 EMSA titration experiments described above (Fig. 1*C*) showed a gradual increase in size of the shifted band in the presence of pre-saturating concentrations of the Csy:F9 complex. This behavior is likely the result of additional molecules of Csy:F9 binding to the DNA as the concentration of the complex increases.

**Fig. 2.**
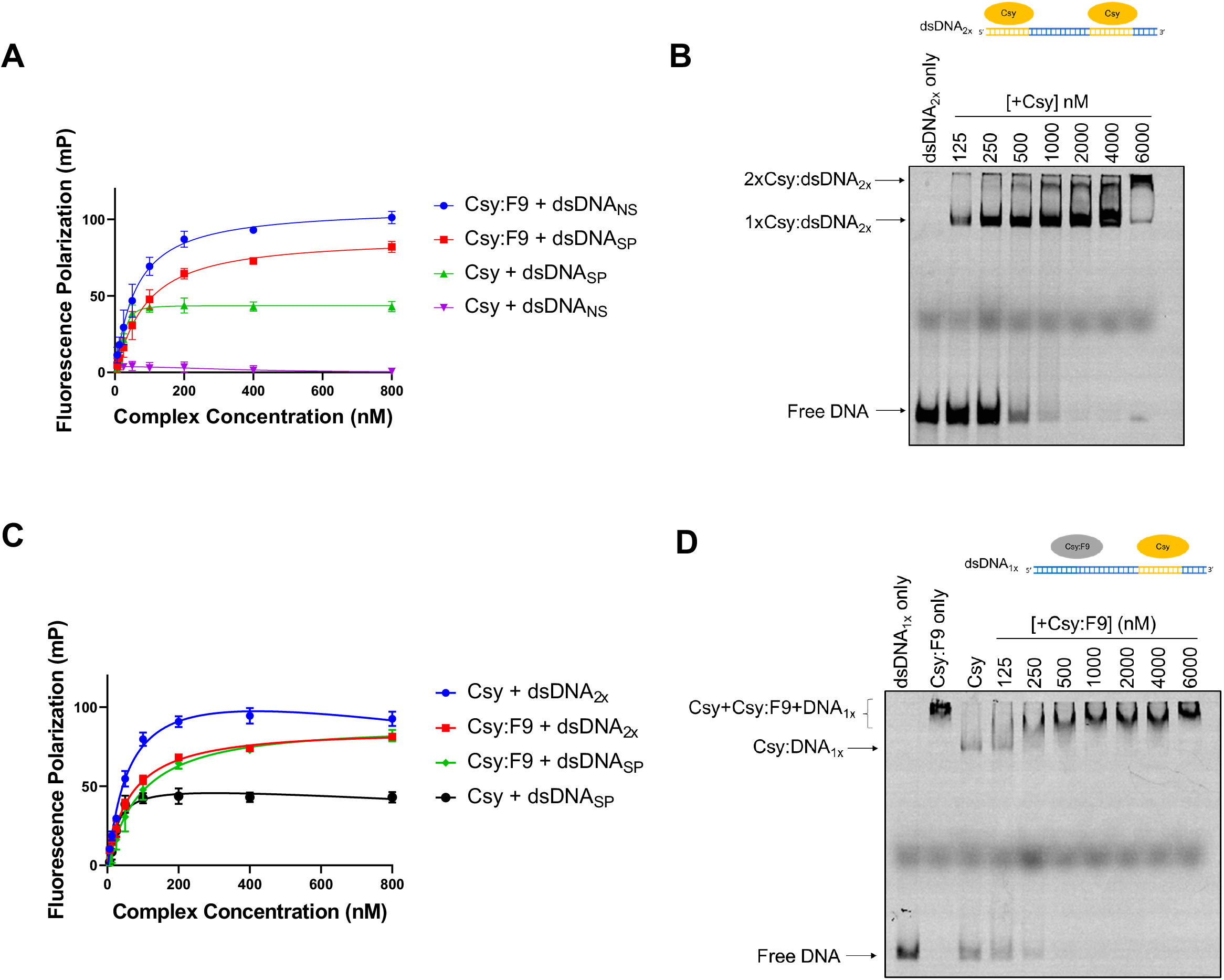
Multiple Csy:F9 can bind to a single piece of dsDNA in a sequence-independent manner. (*A*) The binding of 50 bp Cy5-labeled dsDNA_SP_ and dsDNA_NS_ to Csy and Csy:F9 complexes was assessed by monitoring increase in fluorescence polarization (FP). Increasing concentration of complex were added to a constant concentration of dsDNA ligand. In the legend, the arrow points to the type of DNA being bound by the indicated complex. (*B*) Binding of increasing concentrations of Csy complex to a constant concentration of Cy5-labeled dsDNA_2x_ was monitored on EMSA gels. (*C*) Binding of Cy5-labeled dsDNA_2x_ to increasing concentrations of Csy and Csy:F9 complexes was monitored by FP. FP of Csy and Csy:F9 bound to the 50 bp dsDNA_SP_ from (*A*) are included for comparison. (*D*) Cy5 labeled dsDNA_1x_ was pre-saturated with Csy complex, and then increasing concentrations of Csy:F9 complex were added. DNA-binding was monitored by EMSA. The error bars in (*A*) and (*C*) correspond to standard deviation (SD), *n* = 3.

To directly address the effect of multiple Csy complexes binding to a single DNA molecule, we designed a 60 bp dsDNA target sequence containing two 24 bp complementary binding sites for the Csy complex, each with their own PAMs and including the seed region (henceforth called dsDNA_2X_, Fig. 2*B*). A 24 nt ssDNA molecule complementary to the 5’ end of the crRNA was previously shown to bind strongly to the Csy complex (Bondy-Denomy et al., 2015). EMSAs with dsDNA_2X_ revealed the formation of two distinct bands when mixed with the Csy complex at high concentrations (Fig. 2*B*). The more slowly moving band presumably resulted from the binding of two Csy complexes to one molecule of dsDNA. This band runs with a mobility similar to the band observed when the Csy:F9 complex is mixed with the dsDNA_SP_ or dsDNA_NS_ molecules tested above (Fig. 1*C*). Binding of the Csy complex to dsDNA_2X_ in FP experiments caused a doubling of the FP signal compared to binding of dsDNA_SP_, resulting in a signal level similar to that seen when the Csy:F9 complex binds to dsDNA_SP_, dsDNA_NS_, or dsDNA_2X_ (Fig. 2*A*, 2*C*). A final 60 bp molecule, called dsDNA1X, was synthesized that contained one 24 bp complementary binding site for the Csy complex at one end followed by a random sequence that could act as a non-specific binding site for Csy:F9 (Fig. 2*D*). We first saturated dsDNA1X with the Csy complex, resulting in a single shifted band on the EMSA gel. Subsequent addition of Csy:F9 led to a stepwise slowing of the DNA mobility as the concentration of Csy:F9 was increased (Fig. 2*D*). These data further demonstrate that the slowed mobility of the Csy:F9:DNA complex is the result of multiple complexes binding to a single molecule of DNA.

### Csy:F9 binds dsDNA_NS_ and dsDNA_SP_ at an overlapping site, but ssDNA_SP_ is bound differently

The surprising ability of Csy:F9 to bind dsDNA_NS_ may involve a distinct surface on the Csy complex that is not normally engaged in DNA-binding. We performed competition EMSA experiments to determine whether dsDNA_NS_ and dsDNA_SP_ compete for the same binding site on Csy:F9. The Csy and Csy:F9 complexes were first pre-saturated with FAM-labeled dsDNA_SP_ and then increasing concentrations of Cy5-labeled dsDNA_NS_ were added. In reactions with Csy complex alone, addition of dsDNA_NS_ even at a 4-fold excess caused no reduction in binding to dsDNA_SP_ (Fig. 3*A*, left panel). By contrast, addition of dsDNA_NS_ to Csy:F9 led to increased levels of free dsDNA_SP_, even when the two types of DNA were present at equal concentrations (Fig. 3*A*, left panel). Moreover, most of the dsDNA_SP_ was displaced at a dsDNA_NS_:dsDNA_SP_ ratio of 4:1 (Fig. 3*A*, left panel). Viewing the binding of dsDNA_NS_ by illuminating the same gel with light at 635 nm (absorption wavelength for Cy5) showed that no dsDNA_NS_ was bound to the Csy complex alone but Csy:F9 bound robustly to dsDNA_NS_ even at its lowest concentration (Fig. 3*A*, right panel). Our observation that dsDNA_NS_ readily competes dsDNA_SP_ off of Csy:F9, implies that dsDNA_NS_ and dsDNA_SP_ bind to an overlapping site.

**Fig. 3.**
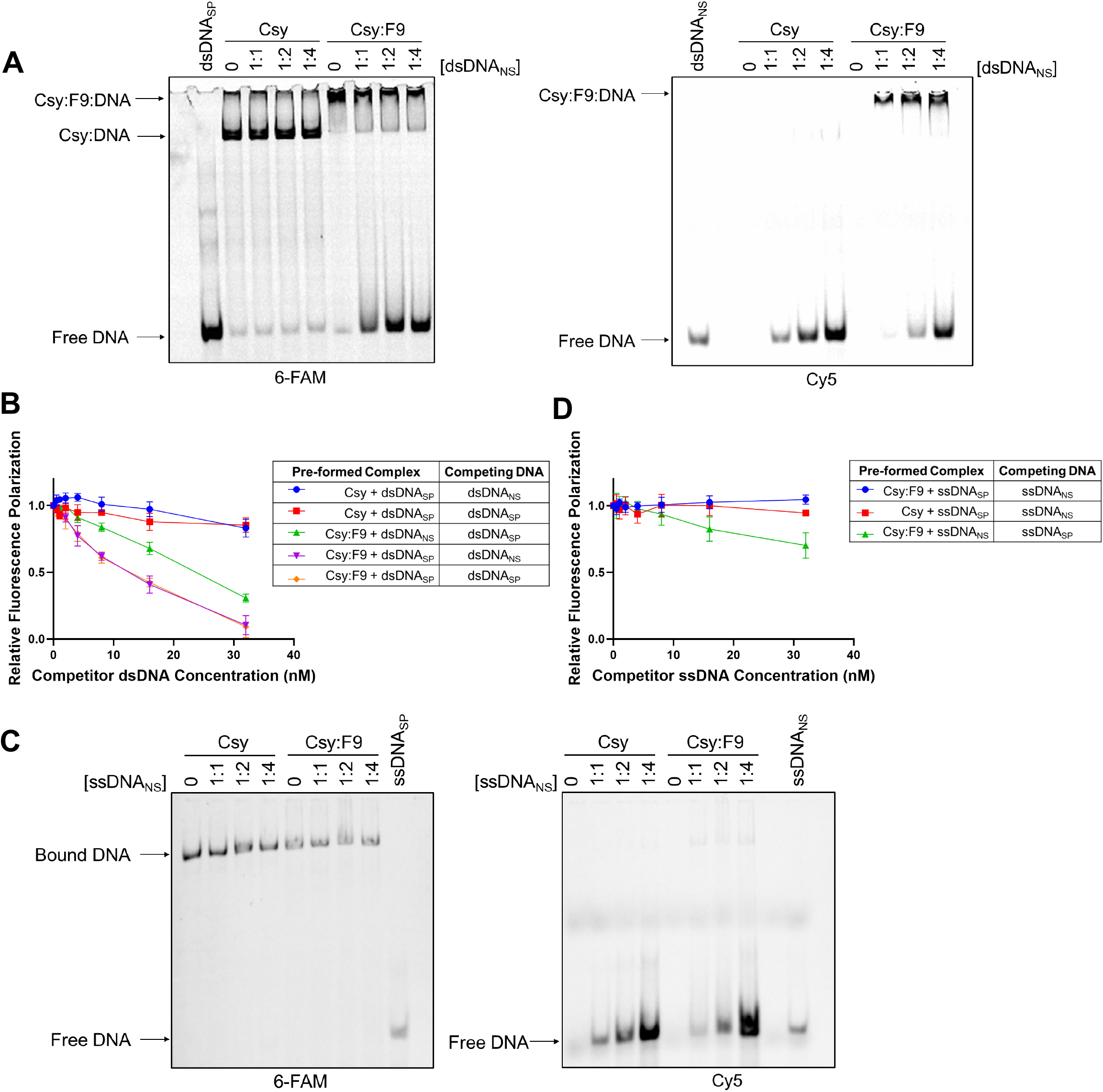
DNA binding competition assays. (*A*) Csy or Csy:F9 complexes (1000 nM) were presaturated with 300 nM 6-FaM labeled dsDNA_SP_, then increasing concentrations of Cy5-labeled dsDNA_NS_ were added as competitor. DNA-binding was monitored by EMSA. The ratio of dsDNA_SP_ to dsDNA_NS_ is shown over each lane. The same gel was irradiated at 473 nm to visualize the 6-FAM-labeled DNA (left panel) and at 673 nm to visualize the Cy5-labeled DNA (right panel). (*B*) Csy or Csy:F9 complexes were pre-mixed with Cy5-dsDNA at 4 nM then competed with increasing concentrations of unlabeled dsDNA as indicated on the x-axis. Relative fluorescence polarization (y-axis) is the FP value at each competitor concentration divided by the value observed in the absence of competitor DNA. (*C*) These competition EMSA experiments are similar to (*A*) except that complexes were pre-saturated with 6-FAM-labeled ssDNA_SP_ and the competitor was Cy5-labeled ssDNA_NS_. (*D*) Competition experiments are similar to (*B*) except that experiments were performed with specific or non-specific ssDNA. The error bars in (*B*) and (*D*) correspond to SD, *n* = 3.

Binding competition experiments were also conducted using FP. Csy or Csy:F9 complexes were pre-saturated with labeled DNA and then challenged with increasing concentrations of non-labeled DNA. If the added non-labeled DNA competed for the same binding site as the labeled DNA, then a decrease in FP signal was expected as the labeled DNA would be competed off of the complex. In the case of the Csy:F9 complex, it can be seen that dsDNA_NS_ and dsDNA_SP_ competed with each other readily and to a similar degree regardless of whether the complex was pre-saturated with dsDNA_SP_ or dsDNA_NS_ (Fig. 3*B*). By contrast, binding of the Csy complex to dsDNA_SP_ was competed negligibly by dsDNA_NS_ even at an 8-fold excess. Notably, the addition of excess levels dsDNA_SP_ to the pre-formed Csy:dsDNA_SP_ complex resulted in little loss of FP signal (Fig. 3*B*), implying that the off-rate of dsDNA_SP_ from Csy is considerably longer than the 30 min incubation period after competitor DNA was added.

To directly address the kinetics of dsDNA binding to Csy:F9, increasing concentrations of unlabeled dsDNA_NS_ were added to a complex pre-saturated with labeled dsDNA_SP_ and the dissociation of the labeled DNA was monitored over time using FP (Fig. S4*A*). Due to the experimental set-up, we were unable to measure time points shorter that 5 min. It can be seen that little change in signal was observed between the 5 min and 30 min time points, indicating that the dsDNA_SP_ was completely dissociated within 5 min. The same dissociation kinetics were observed when the Csy:F9 complex was saturated with labeled dsDNA_NS_ and competed with unlabeled dsDNA_SP_. By contrast, dsDNA_SP_ bound to the Csy complex alone was not competed off at all by dsDNA_NS_ after 30 min (Supp Fig. 4*C*). Overall, these experiments show that bound dsDNA dissociates quickly from Csy:F9 complex, while dsDNA bound specifically to the Csy complex dissociates slowly.

Competition EMSA experiments conducted with ssDNA presented a different picture. While dsDNA_SP_ and dsDNA_NS_ readily competed for an overlapping site on Csy:F9, ssDNA_NS_ was unable to compete ssDNA_SP_ off of the Csy:F9 complex (Fig. 3*C*). This result was similar to that obtained in testing the Csy complex where ssDNA_SP_ was also not competed by ssDNA_NS_ (Fig. 3*C*). Competition experiments monitored by FP corroborated the EMSA results, showing that ssDNA_SP_ binding to Csy:F9 or Csy alone was not competed off by ssDNA_NS_, but ssDNA_NS_ was competed off by ssDNA_SP_ to some extent (Fig. 3*D*). The ability of ssDNA_SP_ to only partially displace ssDNA_NS_ suggests that these molecules are not binding to completely overlapping sites. These data indicate that in contrast to the case with dsDNA, Csy:F9 binds ssDNA_SP_ considerably more strongly than ssDNA_NS_. Using EMSAs, we also observed minimal displacement of pre-saturated ssDNA_SP_ when competed with increasing concentrations of dsDNA_SP_ from the Csy:F9 complex (Fig. S5). Overall, these data demonstrate that ssDNA_SP_ is unique in its stronger binding to Csy:F9 as compared to any other of the DNA molecules tested, implying that Csy:F9 retains the ability to distinguish specific from non-specific ssDNA.

### Cas8f is involved in the binding of dsDNA_NS_ by Csy:F9

The structure of AcrIF9 shows interaction with both Cas7f and Cas8f (K. Zhang et al., 2020), a critical subunit responsible for the initial nonspecific scanning of DNA, PAM recognition, and subsequent strand separation (Chowdhury et al., 2017; Guo et al., 2017). We hypothesized that this subunit may play a role in the unusual properties of Csy:F9. AcrIF2 is a Cas8f-binding anti-CRISPR that can bind to Csy in the presence of AcrIF9 (Fig. S2). The Csy:F2 complex displayed greatly reduced dsDNA_SP_ binding as has been previously shown (Bondy-Denomy et al., 2015) and did not bind to dsDNA_NS_ (Fig. 4*A*). Strikingly, addition of AcrIF2 also drastically reduced the ability of Csy:F9 to bind both dsDNA_SP_ and dsDNA_NS_. This result implied that Cas8f is playing a role in the non-specific dsDNA binding activity of Csy:F9 even though AcrIF9 binds to Cas7f.

**Fig. 4.**
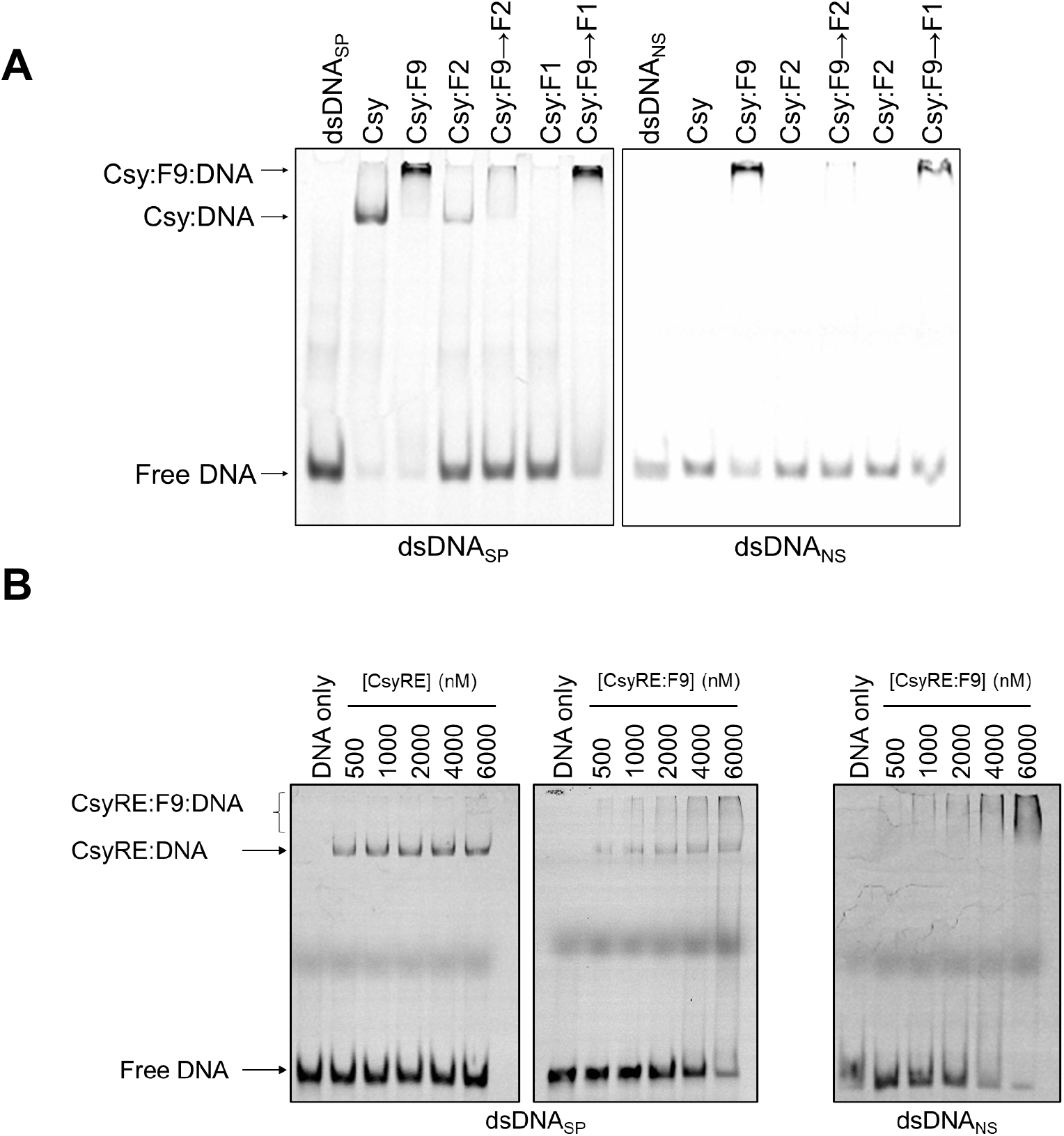
The Cas8f subunit is involved in the Csy:F9 non-specific DNA interaction. (*A*) The Csy complex was incubated with AcrIF9, AcrIF2, AcrIF1, or combinations thereof prior to incubation with 50 bp fragments of FAM-6-labeled dsDNA_SP_ (left) or Cy5-labeled dsDNA_NS_ (right). In cases, where two Acrs were added to one reaction, the Acr added second is indicated by (→). DNA binding activity was assayed by EMSA. (*B*) EMSA gels were used to assess the DNA binding activity of the CsyRE mutant complex in presence (right two panel) or absence (left panel) of AcrIF9. Constant concentrations of dsDNA_SP_ (left 2 panels) or dsDNA_NS_ (right panel) were mixed with increasing concentrations of CysRE or CysRE:F9 complex.

After Cas8f recognizes the PAM, this subunit destabilizes the DNA duplex, leading to base pairing of one DNA strand to the crRNA and to the formation a single-stranded loop, referred to as the R-loop, by the other strand. During this process of PAM recognition, Cas8f undergoes a helical rotation to expose a positively charged channel that binds and stabilizes the R-loop (Rollins et al., 2019). The R-loop binding channel is not sequence specific, as it must bind to a DNA sequence that matches any spacer. To address the possibility that a portion of the non-specific R-loop binding channel might be involved in the non-specific DNA binding activity of Acr:F9, we constructed a mutant Cas8f subunit in which 8 Arg residues lining the R-loop binding channel were replaced with Glu (R207E/R219E/R258E/R259E/R293E/R299E/R302E/R306E). The Csy complex including this Cas8f mutant, which we refer to as CsyRE, displayed reduced binding activity to dsDNA_SP_ as would be expected since this mutant is impaired in its ability to maintain strand separation (Fig. 4*B*). Interestingly, the CsyRE:F9 complex also displayed markedly reduced binding of both dsDNA_SP_ and dsDNA_NS_ (Fig. 4*B*) as compared to the wild-type Csy:F9 complex (Fig. 1*C*). This result further implicates Cas8f in the non-specific DNA binding activity of Csy:F9.

### Non-Specific DNA binding is a conserved property of Csy:F9 complexes

The unusual properties of the Csy:F9 complex led us to question whether this was a conserved feature of this anti-CRISPR or an unusual coincidence occurring with one particular combination of AcrIF9 homologue and Csy complex. To address this issue, we tested a homologue of AcrIF9 from *A. actinomycetemcomitans* that is only 33% identical to the homologue from *P. penneri* used in our other experiments, but still strongly inhibits the type I-F CRISPR-Cas system of *P. aeruginosa* (Fig. S1*B*). Assessment of the DNA binding activity of a Csy:F9_*Aac*_ complex by EMSA showed that it also bound to dsDNA_NS_ in a similar manner as the Csy:F9_*Ppe*_ complex tested above, including the smeared supershifted band (Fig. S6).

To further highlight the universality of Csy:F9 behavior, we assessed the activity of the AcrIF9 homologue from *V. parahaemolyticus* (72% identical to F9_*Ppe*_) against the type I-F CRISPR-Cas system from *Shewanella baltica* (*Sba*,Cas7f 57% identical to Cas7f of *P. aeruginosa*). Using an *in vivo* plasmid transformation assay, it was shown that the *S. baltica* system was completely inhibited by AcrIF9_*Vpa*_, as strains expressing the anti-CRISPR protein and the complex exhibited transformation efficiencies similar to those carrying a deficient nuclease (Cas2-3 HD domain mutant, Fig. S7*A*). The DNA binding properties of Csy_*Sba*_:F9_*Vpa*_ were investigated using Bio-layer Interferometry (BLI). In this approach, biotinylated dsDNA target oligonucleotides were immobilized on a streptavidin-coated bio-sensor. The Csy*Sba* complex was then flowed into the cell and the shift in wavelength of reflected light was measured over time to determine the on-rate (*k*_on_) of the reaction. The off-rate (*k*_off_) was determined by flowing buffer into the cell after the binding reaction reached equilibrium. The BLI experiments showed that Csy_*Sba*_ bound dsDNA_sp_, but showed weak binding to dsDNA_NS_, similar to the binding of AcrIF9_*Vpa*_ alone (Fig. 5*A*, *B*; Fig. S7*B*). By contrast, the Csy_*Sba*_:F9_*Vpa*_ complex was able to bind both dsDNA_SP_ and dsDNA_NS_ (Fig. 5*A*, *B*; Fig. S7*B*). For the specific target, the Csy_*Sba*_:F9_*Vpa*_ complex bound with 14-fold faster *k*_on_ than the Csy_*Sba*_ complex alone (130050 ± 3674.5 M^-1^s^-1^ vs. 9023 ± 860.5 M^-1^s^-1^), and also displayed an 8-fold faster *k*_off_ (0.01805 ± 0.0003 s^-1^ vs. 0.0022675 ± 0.0016 s^-1^).

**Fig. 5.**
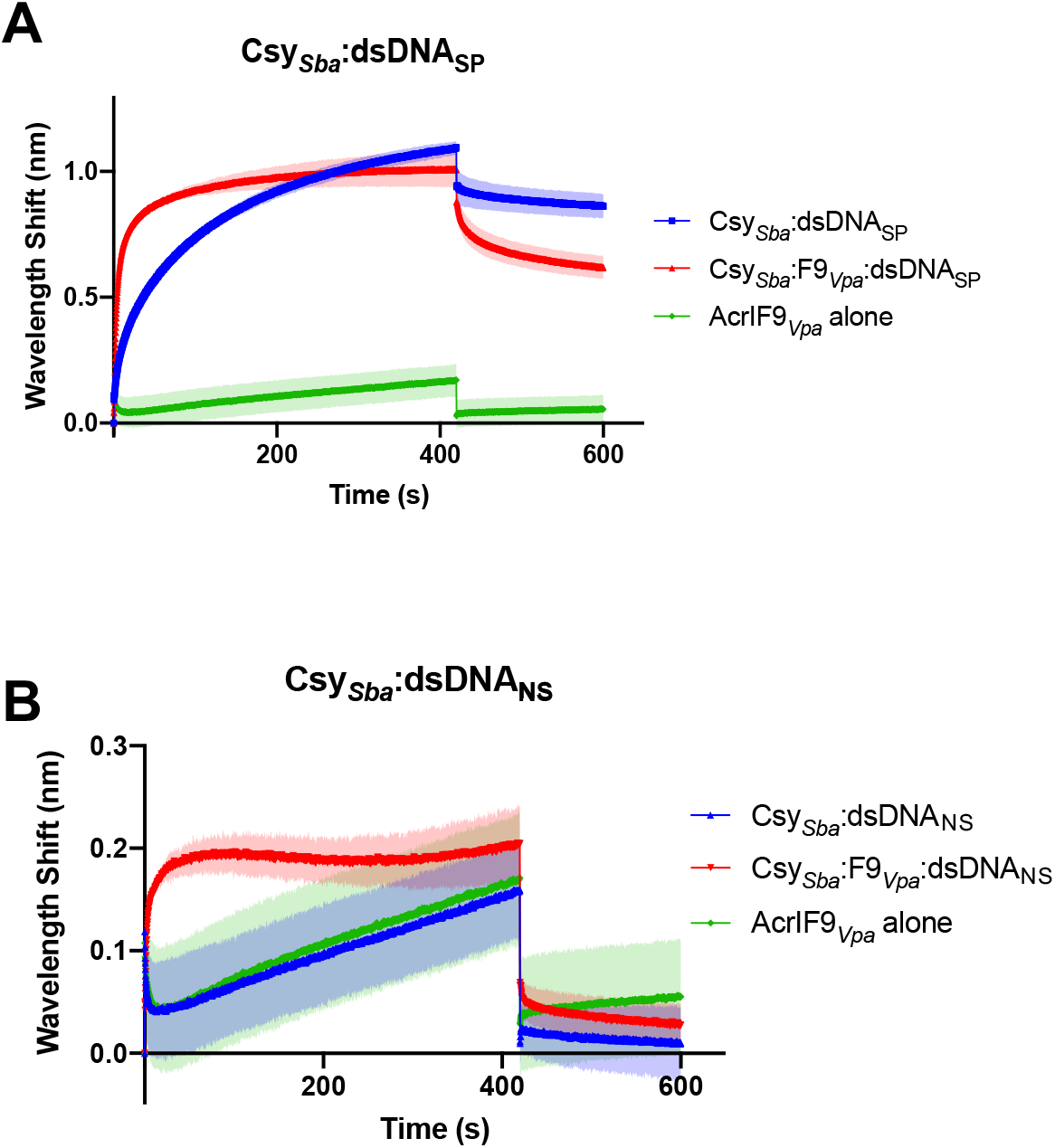
Binding of Type I-F Csy complex from *S. baltica* (Csy_*Sba*_) to dsDNA measured by Bio-Layer Interferometry (BLI) (*A*) The wavelength shift (nm) generated by the binding of Csy_*Sba*_ complex to immobilized dsDNA_SP_ fragments was measured. AcrIF9_*Vpa*_ was mixed with the Csy complex at a 1:1 molar ratio for 10 min at room temperature prior to the initiation of the BLI assay. The shift elicited by AcrIF9_*Ppa*_ binding to dsDNA_SP_ is shown as a control (AcrIF9_*Ppa*_ alone). Assays were performed in duplicates, the lighter outline represents the standard error from the mean. (*B*) The same experiments as described in (*A*) were performed using dsDNA_NS_.

### AcrIF1 and AcrIF9 have different effects on Csy complex stability

The cryo-EM structure of AcrIF9 bound to the Csy complex shows that its binding site overlaps very closely with that of AcrIF1(Fig. 6*A*) (Chowdhury et al., 2017; Guo et al., 2017), and the overall conformation of the Csy:F9 and Csy:F1 complexes are very similar. However, in contrast to the Csy:F9 complex, the Csy:F1 complex displays no non-specific DNA binding activity and is also completely unable to bind specific dsDNA or ssDNA (Bondy-Denomy et al., 2015). These activity differences may result from differences in the conformational flexibility of these two complexes. As a means to interrogate the overall flexibility of the Csy complex alone and in combination with anti-CRISPRs, we treated these complexes with the protease Thermolysin at 55 °C. We chose this treatment because appreciable digestion of Cas7f was not observed at lower temperatures using Trypsin. Thermolysin treatment of the unbound Csy complex resulted in rapid degradation of Cas8f and Cas5f and very little disappearance of the band corresponding to Cas6f (Fig. S8*A*). The band corresponding to Cas7f exhibited intermediate behavior, decreasing steadily over the 5, 10, and 15 min timepoints until only a faint band remained at 30 min. While the degradation pattern of Csy:F9 was very similar to the unbound complex (Fig. S8*B*), the Csy:F1 complex was considerably more resistant to proteolysis (Fig. S8*C*). In particular, the band corresponding to Cas7f was more pronounced at every time point (Fig. 6*B*) and many more bands are visible (Fig S8C). We presume that these extra bands are partial degradation products of Cas7f. A band at the size of AcrIF1 was also visible even after an hour whereas no band corresponding to AcrIF9 was visible even at 5 min. These data indicate that the interaction of AcrIF1 with Cas7f imparts substantial conformational stability, resulting in increased resistance to proteolysis. This stabilization is a property only of AcrIF1 binding even though AcrIF9 binds to the same interface on Cas7f.

**Fig. 6.**
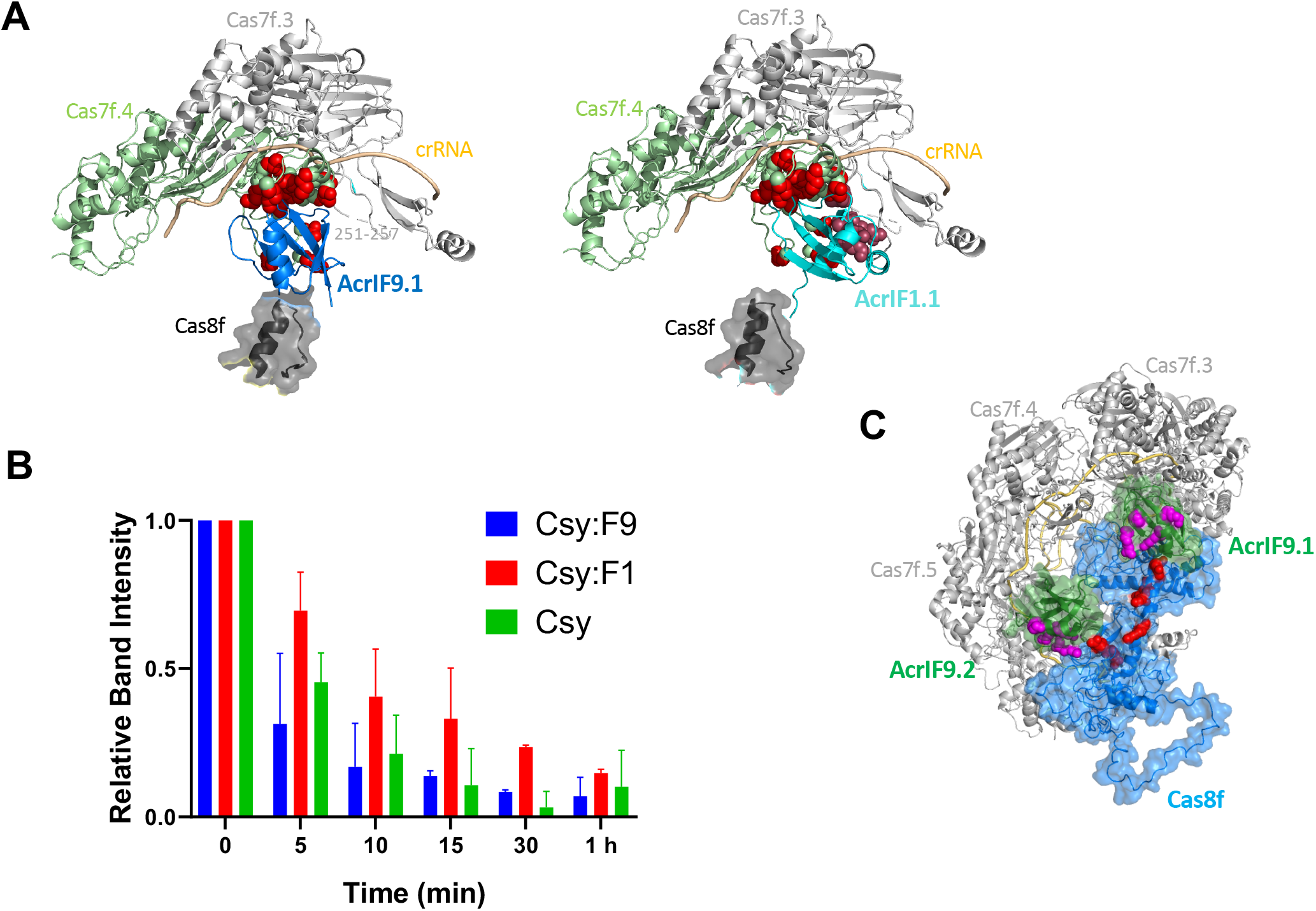
Features of the interactions of AcrIF9 and AcrIF1 with the Csy Complex. (*A*) The interaction interfaces of AcrIF9.1 (left) and AcrIF1.1 (right) with the Csy complexes are shown. Residue side chains in Cas7f.4 that interact with both AcrIF9.1 and AcrIF1.1 are shown in red. When bound to AcrIF9.1, a loop in Cas7f.4 (residues 251-257) indicated in the left panel is disordered. When AcrIF1.1 is bound this loop is ordered and forms intersubunit bridging interactions. Cas7f.4 residues in this interface from the AcrIF1 bound Csy complex (PDB ID 5UZ9) are shown in dark red in the right panel. These figures were made by overlaying the Cas7f subunits of the AcrIF1 bound and AcrIF9 bound (PDB ID 6VQV) structures. These subuntits overlay with less than 2 Å root mean square deviation. (*B*) Csy, Csy:F9, and Csy:F1 complexes were treated with the protease, thermolysin, at 55 °C for various times. Coomassie Blue stained SDS polyacrylamide electrophoresis gels were analyzed to monitor the degradation of Csy subunits over time (Fig. S7). The intensity of the band corresponding to Cas7f at each time point is shown. The Cas7f levels at each timepoint were normalized to that of Cas6f, which was not detectably degraded during this experiment. Cas7f levels were also normalize to the amount of Cas7 present at the 0 time point. Error bars represent SD. Three replicates were analyzed. (*C*) A positive charged surface comprised of AcrIF9 and Cas8 residues is shown. Conserved positively charged residues in AcrIF9 are shown in magenta (Fig. S1). Positively charged residues in Cas8f (<u>R207</u>, <u>K216</u>, <u>R219</u>, <u>R224</u>, <u>R258</u>, <u>R259</u>; underlined residues were substituted with Glu in the CsyRE complex) are shown in red.

## DISCUSSION

A striking feature of anti-CRISPRs is their diversity in sequence, structure, and mechanism of action. Here, we describe the activity of AcrIF9, the first example of an Acr that induces a CRISPR-Cas system to bind dsDNA independent of sequence complementarity or PAM. Binding of AcrIF9 to the Csy complex renders it incapable of distinguishing dsDNA_SP_ from dsDNA_NS_, binding each with similar apparent affinity. Furthermore, the strength of the non-specific DNA-binding induced by AcrIF9 is similar to the specific DNA-binding affinity of the Csy complex on its own, a remarkable property given that non-specific DNA binding activity mediated by Csy complex is so weak as to be undetectable in our assays.

Our *in vitro* DNA binding experiments clearly show that dsDNA_SP_ and dsDNA_NS_ compete equally for the same site on the Csy:F9 complex, indicating that specific hydrogen bonding with the crRNA is not contributing to the interaction (Fig. 3*A*, *B*). The non-specific nature of Csy:F9 DNA binding is also supported by the ability of multiple Csy:F9 complexes to bind one molecule of DNA resulting in the supershifted complexes seen in EMSA gels and the increased signal seen in the FP assays (Fig. 1*D*, 2*A*). Although the affinity of Csy:F9 for non-specific DNA appears to be similar to that of the Csy complex alone for specific DNA, Csy:F9 binds to DNA much more quickly (Fig. 5*A*, *B*), but also dissociates more quickly. These properties are consistent with a reaction driven primarily by electrostatics as would be expected for non-specific DNA binding. The smeared appearance the Csy:F9:DNA complexes in EMSA gels is likely a result of the dissociation of these complexes during electrophoresis. The abrogation of the dsDNA binding activity of Csy:F9 by AcrIF2 or the Cas8f amino acid substitutions (Fig. 4) indicate that Cas8f forms at least part of the non-specific DNA binding interface. The structure of the Csy:F9 complex shows that some of the positively charged Cas8f residues substituted with Glu in the CsyRE complex lie close to positively charged residues in AcrIF9 (Fig. 6*C*). The juxtaposition of positively charged residues in AcrIF9 and Cas8f may create a unique surface for non-specific DNA binding. Notably, a loop of Cas8f between residues 222 and 230 is positioned within less than 3 Å of the AcrIF9.1 molecule (Fig. 6*A*). This interaction between AcrIF9 and Cas8f, which is not seen in the Csy:F1 complex, may stabilize a conformation required for non-specific DNA binding. A recently published cryoEM structure of Csy:F9 bound to DNA indicates a direct involvement of the positively charged residues of AcrIF9 in binding (Hirschi et al., 2020). However, further mutagenesis studies coupled with biochemical assays are still required to corroborate this structural model.

Given the distinct effects of AcrIF1 and AcrIF9 on the activity of the Csy complex, it is surprising that these anti-CRISPRs bind to the Csy complex in such a similar manner. The interactions of both anti-CRISPRs are primarily with one Cas7f subunit to a region known as the thumb (residues 71-95) and to a loop between residues 250 and 261 (Fig. 6*A*). They contact mostly the same residues in these regions even though their sequences and structures are different. A potentially important distinction between these anti-CRISPRs is that AcrIF1 contacts residues 251-257 in the neighboring Cas7f subunit forming an additional interface of 275 Å^2^ (Fig. 6*A*). This same region was unresolved in the Csy:F9 structure, and AcrIF9 makes minimal contact with adjacent Cas7f subunits. The bridging contacts between Cas7f subunits mediated by AcrIF1 may explain the increased resistance of Cas7f to proteolysis when bound to this anti-CRISPR as opposed to AcrIF9 (Fig. 6*B*). AcrIF1 and AcrIF9 appear to sterically occlude the crRNA in a similar manner and both would be expected to block all DNA binding. Consistent with the structure, AcrIF1 does completely inhibit binding of both ssDNA and dsDNA. However, the ability Csy:F9 to distinguish ssDNA_SP_ from ssDNA_NS_ (Fig. 3*C*, *D*) implies that complementary base pairing between crRNA and ssDNA can still occur in this complex. This may due to differences in conformational flexibility between Csy:F9 and Csy:F1 that are not evident from the cryo-EM structure, but may be reflected in the proteolysis experiment.

The non-specific DNA binding activity mediated by AcrIF9 may play an important biological role, or it could be an adventitious *in vitro* curiosity. Our observation that the highly diverged AcrIF9_*Aac*_ elicits non-specific dsDNA binding by the *Pae* Csy complex and AcrIF9_*Vpa*_ does the same when bound to the *Sba* Csy complex demonstrates that this is a conserved property. Furthermore, an alignment of seven diverse AcrIF9 homologues reveals 4 completely conserved positively charged positions (Fig. S1*A*). The residues at these positions are highly exposed the Csy:F9 structure. They also interact with DNA in the structure of the Csy:F9:DNA complex (Hirschi et al., 2020). The 4 other positions in the AcrIF9 alignment that are completely conserved play key functional roles: one (Ala34) is completely buried in the protein core, two (Gln38 and Trp54) are buried in the interface with Cas7f and one (Tyr5) interacts with Cas8f. Substitutions at Gln38 and Tyr5 abrogate AcrIF9 function (Makarova et al., 2020; K. Zhang et al., 2020). Thus, the four surface exposed positively charged positions on AcrIF9 display unusually high conservation equivalent to functionally crucial positions, implying that these residues play a role that has been selected for during evolution. By blocking specific recognition of dsDNA and inducing non-specific DNA binding, AcrIF9 may increase its effectiveness by sequestering Csy complexes in non-productive interactions with bulk DNA within the cell. These non-specific interactions would likely not lead to DNA cleavage because without proper hybridization to the crRNA, the Csy complex is unable to undergo the correct conformational changes required for Cas3 recruitment to the Cas8f subunit (Rollins et al., 2019). Supporting this idea, AcrIF9 blocks *in vitro* cleavage activity mediated by the Csy complex and Cas3 (K. Zhang et al., 2020). Overall, AcrIF9 and its homologues appear to increase their effectiveness by both blocking dsDNA from base pairing with the crRNA to induce Cas3-mediated DNA cleavage, and by diverting the Csy complex away from the target DNA through non-specific DNA binding. The unique nature of AcrIF9 inhibition may partially explain the widespread distribution of members of this Acr family and their ability to block diverse type I-F systems (Pawluk et al., 2016). Interestingly, another very widespread Acr, AcrIIA11, possesses DNA binding activity on its own, and this activity is increased in the presence of Cas9 (Forsberg et al., 2019).

In conclusion, we have shown that AcrIF9 possesses the remarkable ability to convert the normally highly specific dsDNA binding activity of the Csy complex into a completely non-specific binding activity that is, nevertheless, still very strong. The potential to become a strong non-specific DNA binding complex may be inherent in all dsDNA binding CRISPR-Cas systems as they all must form non-specific interactions with the single strand of DNA that is looped out (the R-loop) upon hybridization of the target strand to the crRNA. In the case of Csy:F9 complex, the surface of Cas8f responsible for R-loop binding appears to be involved in the stabilization of non-specific DNA interactions. Given the known conformational flexibility of the Csy complex (Guo et al., 2017), it is possible that AcrIF9 may preferentially stabilize conformations that expose residues involved in non-specific DNA binding. Further investigation of AcrIF9 and other Acrs with unexpected activities will provide unique pathways to increase our understanding of the intricacies of CRISPR-Cas function.

## MATERIALS AND METHODS

### In vivo assay for Acr activity

In vivo assays to detect anti-CRISPR activity were carried out as originally described (Bondy-Denomy et al., 2013). pHERD30T (Qiu, Damron, Mima, Schweizer, & Yu, 2008) derived plasmids were used to express AcrIF9 homologues in *P. aeruginosa* strain UCBPP-PA14 (PA14), which possesses an active type I-F CRISPR-Cas system. Lysates of a CRISPR-Cas sensitive phage (DMS3m) or a CRISPR-Cas insensitive phage (DMS3) were spotted in ten-fold dilutions onto lawns of PA14 transformed with plasmids expressing Acrs of interest. A strain carrying pHERD30T was used as a negative control. Plates were incubated at 30 °C overnight. Homologues to be tested were identified by PSI-BLAST (2 iterations) (Altschul & Koonin, 1998). The protein sequence alignment was constructed and analyzed using Jalview (Waterhouse, Procter, Martin, Clamp, & Barton, 2009).

### Expression and purification of Csy complex and Acrs

The *P. aeruginosa* Csy complex including crRNA was expressed from plasmids in *E. coli* strain BL21(DE3) as previously described (Wiedenheft et al., 2011). Cas7f is tagged with 6xHis. To produce Csy:F9, the constructs expressing Csy complex and crRNA as stated above were coexpressed with pCDF-1b expressing untagged AcrIF9.

Cultures of *E. coli* BL21 (DE3) expressing the protein of interest were grown to an optical density (OD600) of 0.6 and then induced with 1 mM isopropyl-b-D-thiogalactoside (IPTG) for 16h at 16 °C. Cells were collected by centrifugation at 7000 g for 15 min and resuspended in binding buffer (20 mM Tris pH 7.5, 200 mM NaCl, 5 mM imidazole, 1 mM tris (2-carboxyethyl)phosphine (TCEP)). The cells were lysed by sonication and the resulting lysates were centrifuged at 17,000 g for 25 min to remove cell debris. The supernatant was mixed with Ni-NTA beads and incubated for 1 hr at 4 °C. The lysates and the beads were then passed through a column, washed 5 times with wash buffer (20 mM Tris, pH 7.5, 200 mM NaCl, 30 mM imidazole, 5 mM β-mercaptoethanol) and then eluted in buffer containing 300 mM imidazole. Purified protein was dialysed into 20 mM Tris pH 7.5, 200 mM NaCl, 1 mM TCEP) overnight. Affinity-purified proteins were fractionated by size exclusion chromatography (SEC) using a GE Life Sciences Superdex 200 16/600 column. Fractions were collected in 1.5 ml volumes and monitored by optical density at 280 nm. Protein purity was assessed by visualization on Coomassie blue R250 stained SDS–PAGE gels.

### Assessing Acr binding to the Csy complex

Purified 6xHis-tagged Csy complex (1000 nM) was bound to Ni-NTA beads and incubated with excess Acr (5000 nM) for 30 min at 4 °C in binding buffer (20 mM Tris pH 7.5, 200 mM NaCl, 5 mM imidazole, 1 mM TCEP). Competitor Acr was added in equimolar concentration and incubated for 30 min at 4 °C. Bound Csy complex and Acr was collected through centrifugation at 4000 rpm for 2 min to remove unbound Acr. The reaction was then washed three times with wash buffer (20 mM Tris, pH 7.5, 200 mM NaCl, 30 mM imidazole, 5 mM β-mercaptoethanol) with a centrifugation step after each wash. The sample was then eluted in elution buffer (20 mM Tris, pH 7.5, 200 mM NaCl, 300 mM imidazole, 5 mM β-mercaptoethanol). The samples were visualized on Coomassie blue R250 stained SDS–PAGE gels. Each experiment was conducted at least 3 times and the same result was obtained each time. Single representation is shown in Fig. S2.

### Site-directed mutagenesis

Complementary oligonucleotides comprising the codon to be mutated plus 20 nucleotides in both directions were synthesized by Eurofins Genomics. The entire plasmid template was then PCR amplified with the primers containing the mutations using Phusion DNA polymerase.

Subsequently, the template was digested with DpnI and the PCR product was transformed into *E. coli* DH5α. Mutations were confirmed by DNA sequencing.

### DNA binding assays

DNA molecules (sequences shown below) were synthesized (Eurofins Genomics) that contain 32 nucleotides that is either complementary (specific) to the crRNA in the Csy complex or scrambled (nonspecific). The DNA was fluorescently labeled at the 5’ end with either 6-FAM or Cy5. To generate dsDNA, the labeled strand was mixed with an unlabeled complementary strand, heated to 100 °C, and allowed to return slowly to room temperature. DNA binding reactions were conducted in a binding buffer (10 mM HEPES, pH 7.5, 1 mM MgCl2, 20 mM KCl, 1 mM TCEP, and 6% glycerol) at 37 °C for 15 min. A DNA concentration of 100 nM was used in EMSA reactions with Csy or Csy:Acr complexes at 2000 nM. In competitive DNA binding experiments, the Csy complex, or Csy:F9 (1000 nM), were first incubated with 300 nM of DNA at 37 °C for 15 min. Then the competitor DNA was added at increasing concentrations with the following ratios (1:1, 1:2, 1:4) and incubated at 37 °C for another 15 min. For EMSA experiments with competing Acrs, the Csy complex was first incubated with ten-fold excess of one Acr for 1 hour at 4 °C then equimolar amount of the competitor Acr was added and incubated under the same conditions. 100 nm DNA was then incubated with the resulting Acr-bound Csy complex at 37 °C for 15 min. All EMSA reactions were resolved on native 4% or 6% polyacrylamide TBE gels. Gels were visualized with a Typhoon imager at 473 nm (6-FAM) and 635 nm (Cy5). Every EMSA experiment was carried out at least three times with reproducible results. Single representative gels are shown in figures.

For FP assays, Cy5 labeled DNA probes (4 nM) were incubated with purified Csy or Csy:F9 complexes at increasing concentrations (6.25,12.5, 25, 50,100, 200, 400, 800 nM) in a total volume of 50 ul in Greiner Bio-one 96 well black flat-bottom microplates. The samples were mixed with the assay buffer (20 mM Hepes pH 7.5, 50 mM KCl, 5 mM MgCl2, 0.01% Triton X100, 2 mM DTT, 0.1mg/ml bovine gamma globulin) and incubated at 37 °C for 30 min. The plate was then analyzed with a Tecan Microplate Reader Spark at 635 nM. The polarization signal was corrected to the reference (Cy5-DNA only) and the blank (assay buffer only). For competitive assays, the Csy or Csy:F9 complex (100 nM) was first incubated with Cy5-labeled DNA at 4 nM for 30 min at 37 °C and then competed with increasing concentrations of unlabeled DNA (0, 0.5, 1, 2, 4, 8, 16, 32 nM) for 30 min at 37 °C. All FP assays were performed at least 3 times. Average values are plotted with error bars representing standard deviation.

The sequences of the DNA used for DNA binding assays are shown below. Protospacer sequences complementary to the crRNA are in bold and underlined. PAMs are in red. In the 2x and 1x sequences we used a 24 nt protospacer instead of 32 nt in order to keep all dsDNA target sequences used at approximately the same length. We found that the Csy complex binds with similar affinity to a 24 nt protospacer.

50 bp DNA_SP_ target strand:

5’: GAATGACCTA<u>**CAGGTAGACGCGGACATCAAGCCCGCCGTGAA**</u>GGCATGTCAA

50 bp DNA_NS_:

5’:GCGCACCTATTAACCGTTCGCAGAAACCAGTAGTAGTCCAAGCGACATGCAG

DNA_2x_ target strand:

5’: <u>**CGCGGACATCAAGCCCGCCGTGAA**</u>GGCATGTC<u>**CGCGGACATCAAGCCCGCCG TGAA**</u>GGCATGT

DNA_1x_ target strand:

5’: GTAGTAGTCCAACGGCATGTAATGACCTA<u>**CGCGGACATCAAGCCCGCCGTGAA**</u> GGCATGT

*In vivo* activity measurement for Csy_*Sba*_ The efficiency of transformation assay (EOT) was performed as described previously (Pausch et al., 2017). Csy_*Sba*_ was expressed in *E. coli* BL21-AI in the presence and absence of AcrIF9_Vpa_. A spacer targeting the ampicillin resistance cassette of pETDuet-1 was used to determine the EOT. EOT equals to the colony ratio between the colony count of the strain of interest and its corresponding Cas3 HD mutant strain, presented as percentages. Error bars represent the standard error of the mean, three replicates were quantified.

### Bio-layer interferometry (BLI)

The conditions used in the BLI experiments were as described previously (Müller-Esparza, Osorio-Valeriano, Steube, Thanbichler, & Randau, 2020). AcrIF9_Vpa_ and Csy complex were mixed at a molar ratio of 1:1 and incubated for 10 min at room temperature in BLI buffer (0.1 μM BSA and 0.01% Triton X-100). The Csy complex, Csy:F9, or AcrIF9_Vpa_ alone, was tested against 100 nM of either dsDNA_SP_ or dsDNA_NS_. Assays were performed in duplicate on the BLItz platform (FortéBio) using High Precision Streptavidin (SAX) Biosensors (FortéBio).

### Limited proteolysis experiment

Purified Csy or Csy:Acr complexes were diluted to a final concentration of 1000 nM in buffer (20 mM Tris, pH 7.5, 200 mM NaCl, 1 mM TCEP). Samples were incubated at 55 °C with 0.003 mg/ml Thermolysin (Sigma Aldrich). Aliquots were removed at each timepoint into 2x protein sample buffer (62.5 mM Tris pH 6.8, 2.5% SDS, 0.002% Bromophenol Blue, 5% β-mercaptoethanol, 10% glycerol) heated at 100 °C for 5 min to inactivate Thermolysin. Samples were analyzed on Coomassie Blue stained SDS-PAGE gels. Band intensities on the gels were quantitated using ImageJ (Schneider, Rasband, & Eliceiri, 2012). The intensity of Cas7f at each timepoint was normalized to that of Cas6f. The relative band intensity was then determined by taking the ratio between Cas7f at the different time points to that at 0 min.

## Acknowledgements

This work was supported by Canadian Institutes of Health Research grant FDN-15427 (A.R.D.). L.R. was funded by the German Research Foundation (DFG, grant SPP 2141).

---

**Fig S1.**
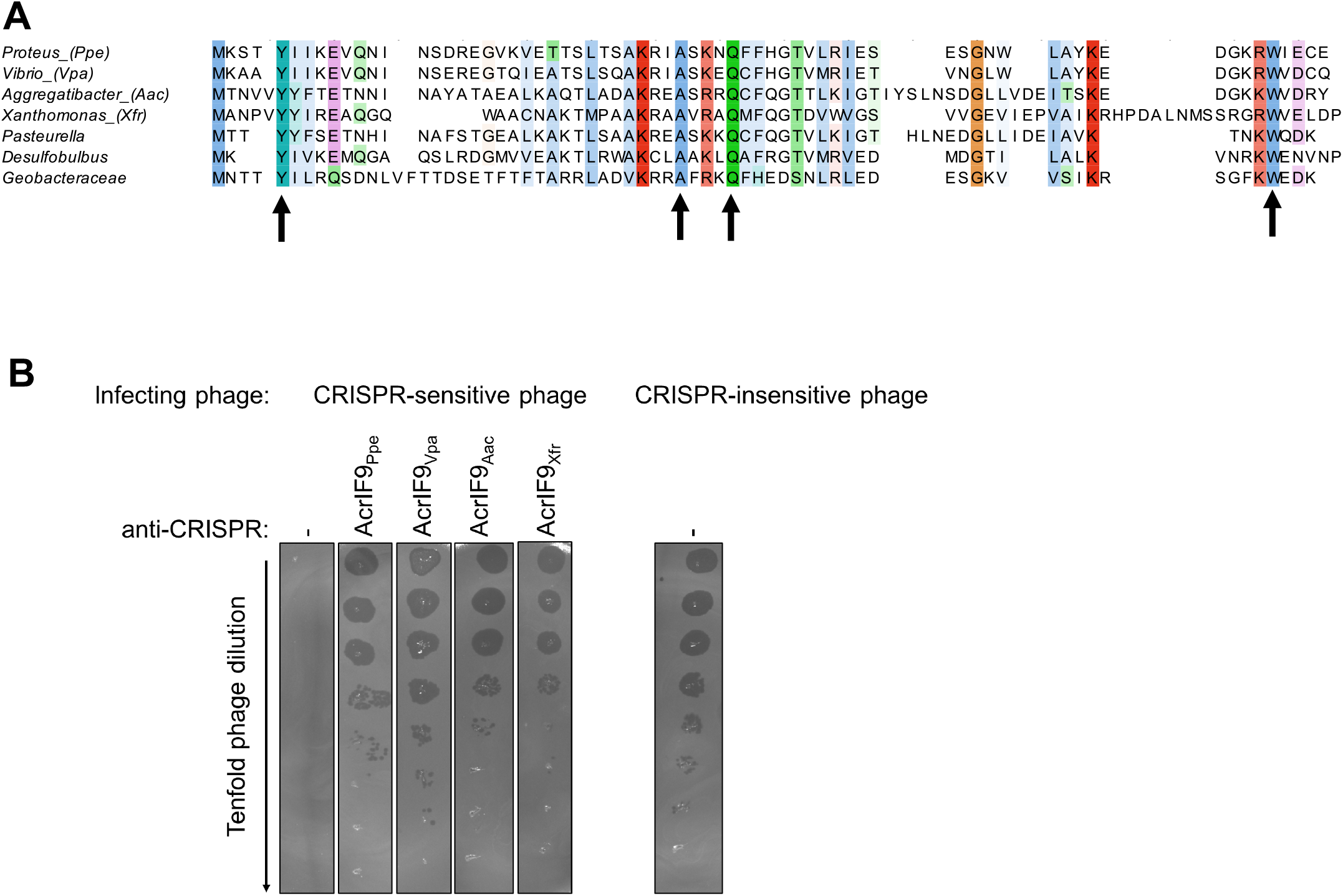
AcrIF9 homologues inhibits the *P. aeruginosa* type I-F CRISPR-Cas system. (*A*) A sequence alignment of 7 diverse members (average pairwise sequence identity = 34%) of the AcrIF9 family. The alignment is colored by conservation according to the ClustalX scheme. The conserved positively charged positions are red and light red. Other completely conserved non-Gly positions are indicated with arrows. The sequences shown are PROPEN_01997 (*Proteus penneri* ATCC 35198), D5E77_24790 (*Vibrio parahaemolyticus*), FXB80_00875 (*Aggregatibacter actinomycetemcomitans*), PD5205_04008 (plasmid) (*Xanthomonas fragariae*), BGK37_12565 (*Pasteurella multocida*), JT06_19085 (*Desulfobulbus* sp. Tol-SR), and A2076_14270 (*Geobacteraceae* GWC2_53_11). (*B*) Dilutions of CRISPR-sensitive (DMS3m) and CRISPR-insensitive bacterium (DMS3) phage lysates were spotted onto lawns of *P. aeruginosa* strain PA14, which has an active type I-F CRISPR-Cas system. PA14 was transformed with plasmids expressing the indicated AcrIF9 homologs. Zones of clearing indicate successful phage infection and inhibition of the CRISPR-Cas system. The negative control is strain PA14 carrying only the expression vector.

**Fig. S2.**
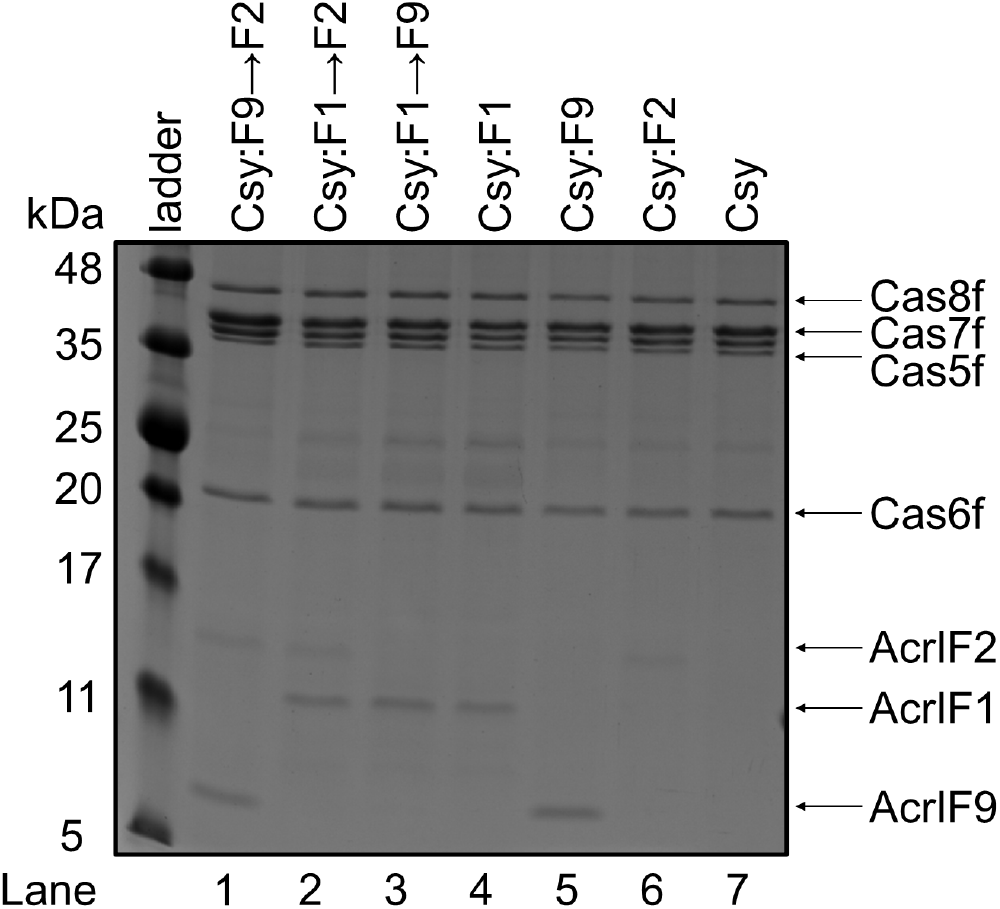
Competition between Acrs for binding to the Csy complex. Untagged Acrs were mixed with the Csy complex containing 6xHis-tagged Cas7f. The samples were then affinity purified using Ni-NTA beads to remove unbound Acr and proteins eluted from the beads were visualize on Coomassie Blue stained SDS-PAGE gels. In competitive binding experiments the Acr following the (→) was added second after preincubation with the first Acr.

**Fig. S3.**
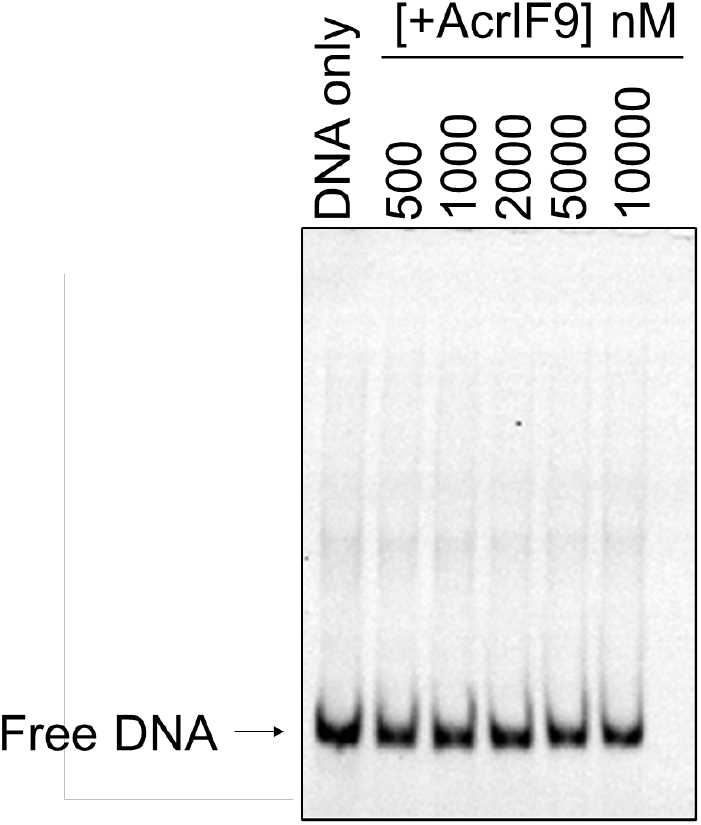
AcrIF9 does not bind dsDNA without the Csy complex. Increasing concentrations of AcrIF9 were mixed with 100 nM of FAM-6 labeled 50 bp dsDNA_SP_fragment. No Csy complex was present. DNA binding was detected by EMSA.

**Fig. S4.**
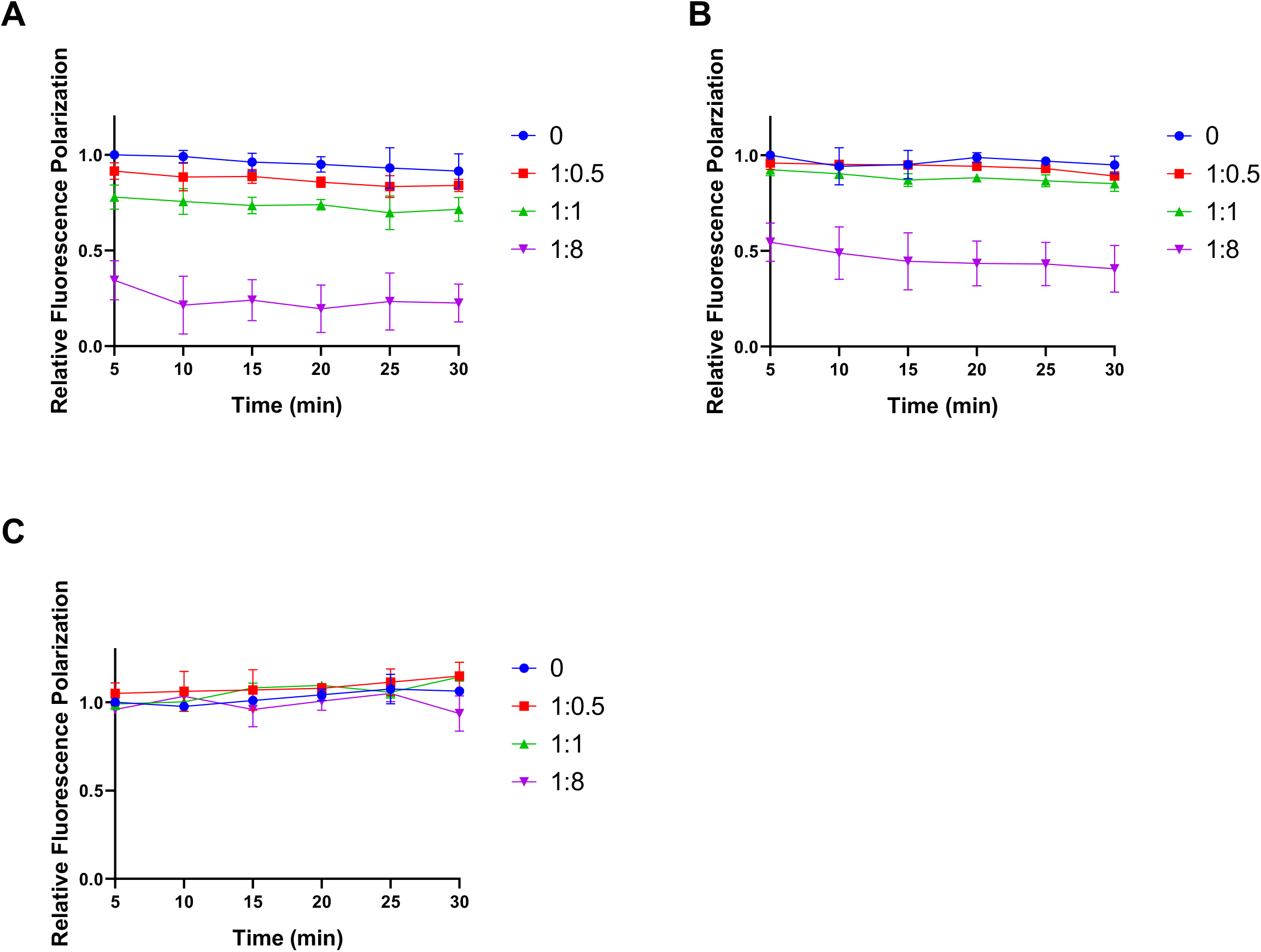
The dsDNA bound by Csy:F9 is rapidly competed off. Csy or Csy:F9 complexes were pre-saturated with Cy5-labeled DNA, then increasing concentrations of unlabeled DNA were added at levels ranging from 0.5 to 8-fold excess over the concentration of the labeled DNA (i.e. ratios of 1:0.5, 1:1, 1:8). FP measurements were taken at various times after addition of the unlabeled DNA. Due to limitations of the apparatus being used, time points shorter than 5 min could not be assayed. Relative fluorescence polarization was calculated using the ratio of the values from each competitor concentration to that without competitor DNA at 5 min. All error bars represent SD, *n* = 3. (*A*) Csy:F9 was pre-saturated with dsDNA_SP_ and competed with unlabled dsDNA_NS_. (*B*) Csy:F9 was pre-saturated with dsDNA_NS_ and competed with unlabeled dsDNA_SP_. (*C*) Csy complex was pre-saturated with dsDNA_SP_ and competed with unlabeled dsDNA_SP_.

**Fig. S5.**
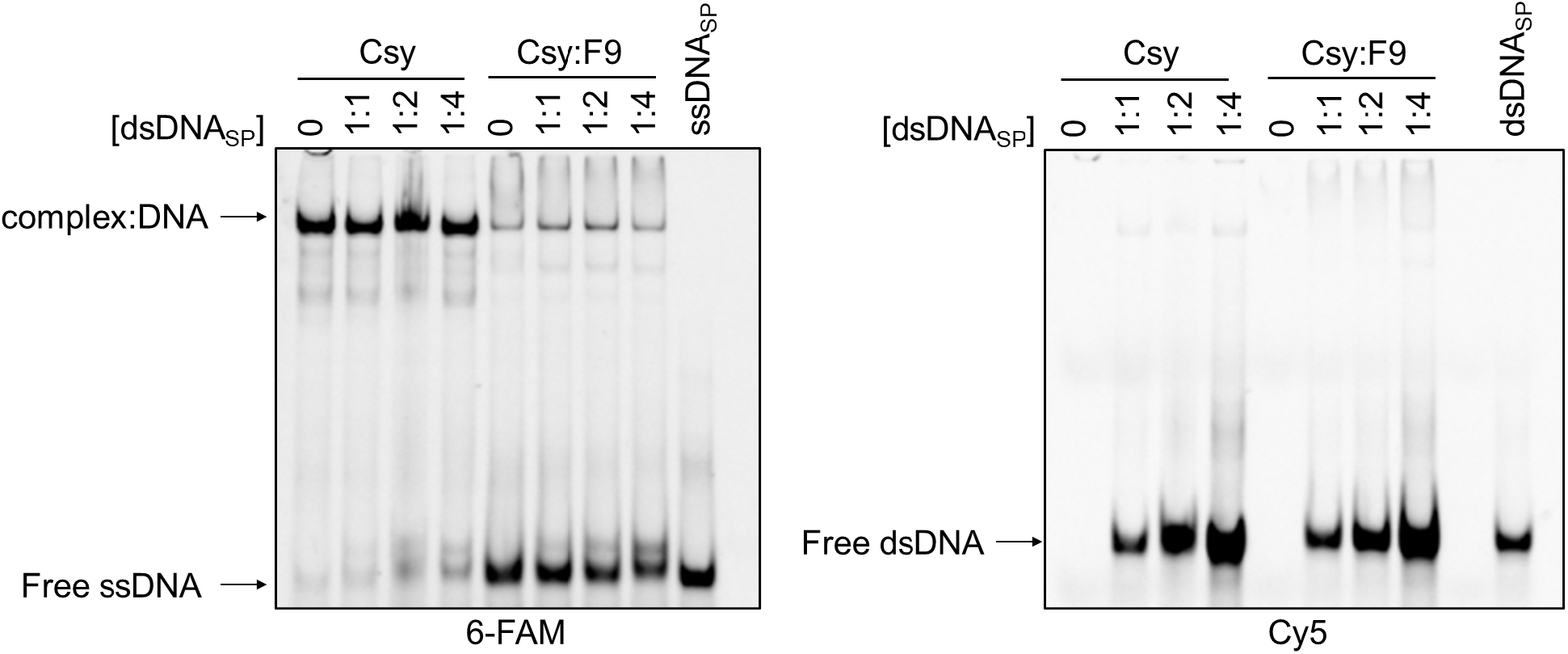
Csy:F9 differentially interacts with dsDNA and ssDNA. Csy or Csy:F9 complexes (2000 nM) were pre-saturated with 1000 nM of 6-FAM labeled ssDNA_SP_, then increasing concentrations of Cy5-labeled dsDNA_SP_ were added as competitor. DNA-binding was monitored by EMSA. The ratio of dsDNA_SP_ to dsDNA_NS_ is shown over each lane. The same gel was irradiated at 473 nm to visualize the 6-FAM-labeled DNA (left panel) and at 673 nm to visualize the Cy5-labeled DNA (right panel).

**Fig. S6.**
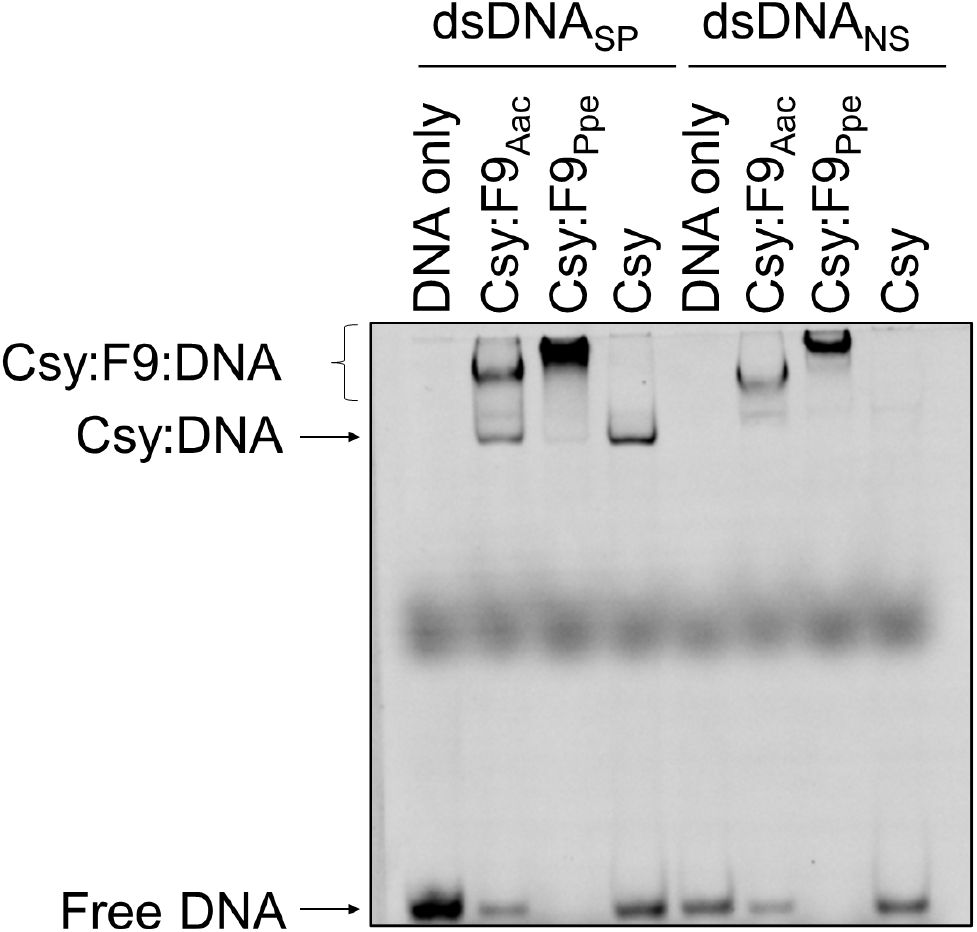
Non-specific DNA binding is a conserved feature of AcrIF9 homologues. EMSA gels showing binding of Csy:F9_Aac_, Csy:F9_Ppe_, and Csy alone to Cy5-labeled dsDNA_SP_ and dsDNA_NS_. These experiments were performed under identical conditions as those shown in Fig. 1*A*.

**Fig. S7.**
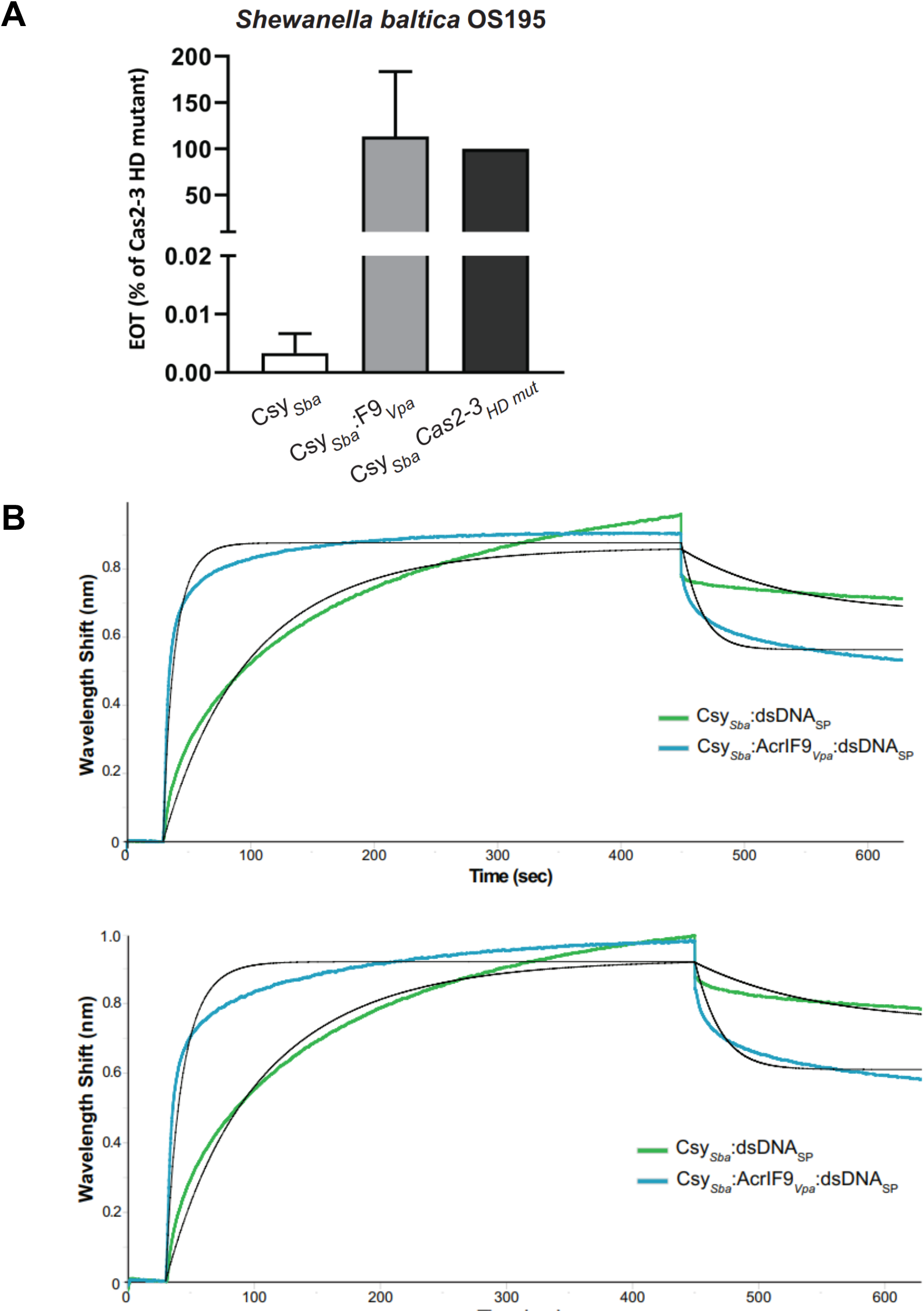
AcrIF9 inhibits the Type I-F system of *S. baltica* (Csy_*Sba*_). (*A*) Efficiency of Transformation assays (EOTs) of the Csy_*Sba*_ complex expressed in *E. coli* BL21-AI were performed. The activity of the effector complexes was tested when expressed alone (Csy_*Sba*_) or coexpressed with AcrIF9_*Vpa*_. EOT is equal to the colony ratio between the strain of interest and its corresponding Cas2-3 HD mutant strain, presented as percentages. Error bars represent the SEM, three replicates were quantified. (*B*) Independent BLI wavelength shifts (nm) generated by the binding Csy_*Sba*_ to a specific dsDNA oligonucleotide, in the presence (blue) or absence (green) of AcrIF9_*Vpa*_ are shown. Each experiment was performed with independently purified protein samples and freshly diluted oligonucleotide stocks. Non-linear regressions obtained by least-square fit by the BLItz Pro Software are shown in black for each curve.

**Fig. S8.**
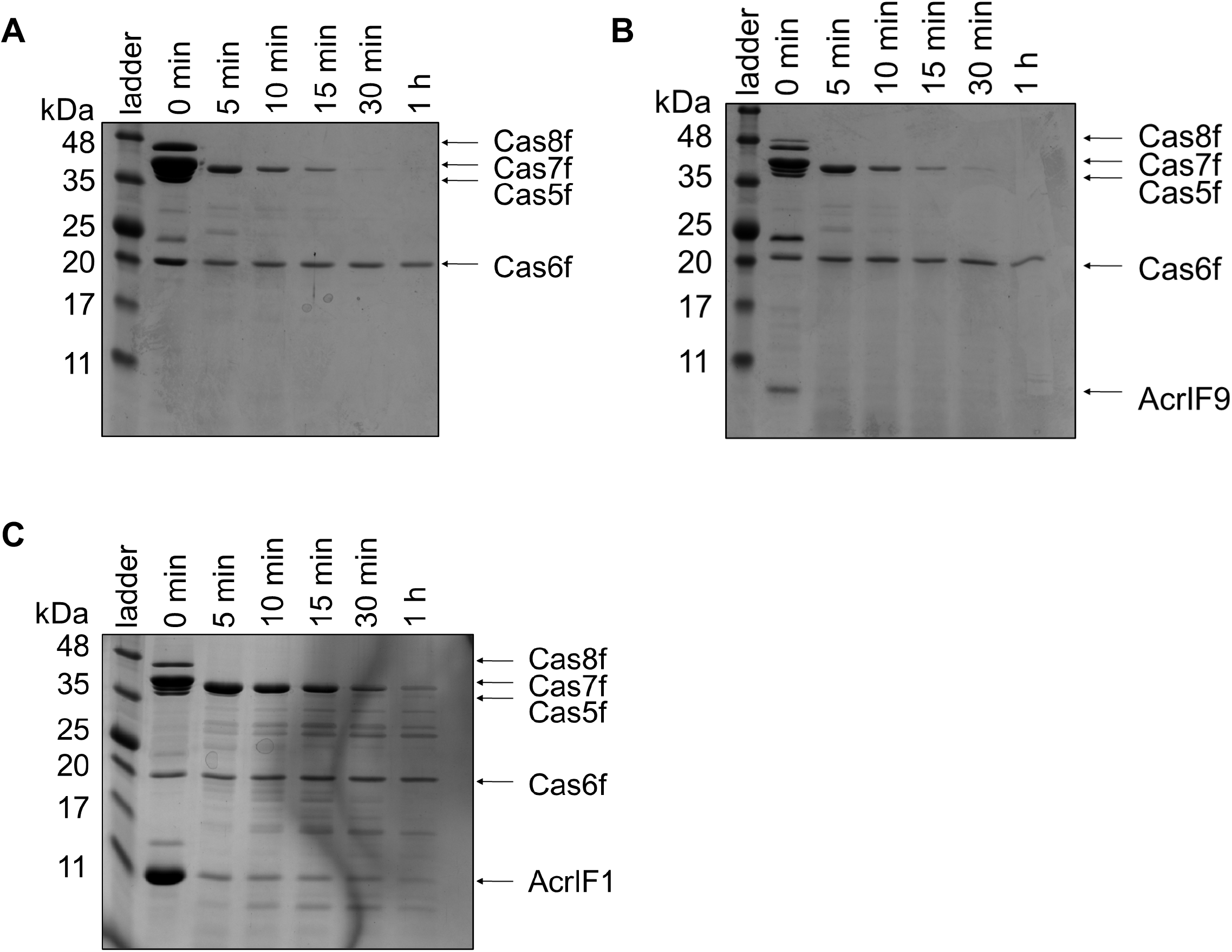
AcrIF1 stabilizes Cas7f against proteolysis, while AcrIF9 does not. Csy (*A*), Csy:F9 (*B*), and Csy:F1 (*C*) complexes were treated with the protease, thermolysin, at 55 °C for various times. Coomassie Blue stained SDS polyacrylamide electrophoresis gels were analyzed to monitor the degradation the Csy subunits over time.

